# A comprehensive profile of circulating RNAs in human serum

**DOI:** 10.1101/186320

**Authors:** Sinan Uğur Umu, Hilde Langseth, Cecilie Bucher-Jonannessen, Bastian Fromm, Andreas Keller, Eckart Meese, Marianne Lauritzen, Magnus Leithaug, Robert Lyle, Trine Rounge

**Affiliations:** Department of Research, Cancer Registry of Norway, Oslo, Norway; Department of Tumor Biology, Institute for Cancer Research, The Norwegian Radium Hospital, Oslo University Hospital, Nydalen, Oslo, Norway; Department of Clinical Bioinformatics, Saarland University, 66041, Saarbruecken, Germany; Department of Human Genetics, Saarland University, 66421, Homburg/Saar, Germany; Department of Medical Genetics, Oslo University Hospital and University of Oslo, Oslo, Norway; PharmaTox Strategic Research Initiative, School of Pharmacy, Faculty of Mathematics and Natural Sciences, University of Oslo, Oslo, Norway

## Abstract

Non-coding RNA (ncRNA) molecules have fundamental roles in cells and many are also stable in body fluids as extracellular RNAs. In this study, we used RNA sequencing (RNA-seq) to investigate the profile of small non-coding RNA (sncRNA) in human serum. We analyzed 10 billion Illumina reads from 477 serum samples, included in the Norwegian population-based Janus Serum Bank (JSB). We found that the core serum RNA repertoire includes 258 micro RNAs (miRNA), 441 piwi-interacting RNAs (piRNA), 411 transfer RNAs (tRNA), 24 small nucleolar RNAs (snoRNA), 125 small nuclear RNAs (snRNA) and 123 miscellaneous RNAs (misc-RNA). We also investigated biological and technical variation in expression, and the results suggest that many RNA molecules identified in serum contain signs of biological variation. They are therefore unlikely to be random degradation by-products. In addition, the presence of specific fragments of tRNA, snoRNA, Vault RNA and Y_RNA indicates protection from degradation. Our results suggest that many circulating RNAs in serum can be potential biomarkers.

## INTRODUCTION

Human serum and plasma contain various classes of RNA molecules (Danielson et al. 2017; Keller et al. 2011; Hornick et al. 2015) such as protein-coding mRNAs (Kim et al. 2017), miRNAs (Inns and James 2015; Leidinger et al. 2013; Rounge et al. 2015; Hornick et al. 2015; Mitchell et al. 2008; Arroyo et al. 2011; Chen et al. 2008), piRNAs (Yuan et al. 2016; Danielson et al. 2017), tRNAs and miscellaneous other ncRNA molecules (Yuan et al. 2016; Danielson et al. 2017). These circulating RNAs are usually packed in extracellular vesicles and have considerable potential as minimally-invasive biomarkers (Yuan et al. 2016; An et al. 2015; Inns and James 2015; Kim et al. 2017; Nolte-’t Hoen et al. 2012; Mitchell et al. 2008), since they are stable and some have been associated with disease phenotypes (Inns and James 2015; Yuan et al. 2016; Nomura 2017; Leidinger et al. 2013; Maierthaler et al. 2017).

miRNAs are the best characterized class of sncRNA molecules. They are approximately 22 nucleotides (nts) in length and regulate cellular gene expression via RNA-RNA antisense binding (Ambros 2004; Chen 2008; Umu and Gardner 2017). They can also be found as circulating RNAs (Inns and James 2015; Leidinger et al. 2013; Rounge et al. 2015; Hornick et al. 2015; Mitchell et al. 2008). Many studies have investigated the biomarker potential of miRNAs (Rounge et al. 2015; Flatmark et al. 2016; Mitchell et al. 2008; Keller et al. 2011; Maierthaler et al. 2017; Mendell and Olson 2012; Inns and James 2015; Leidinger et al. 2013; Arroyo et al. 2011) and their isoforms, isomiRs (Telonis et al. 2017; Morin et al. 2008; Llorens et al. 2013). Small nucleolar RNAs (snoRNAs) are another well-known member of sncRNA molecules. They play a crucial role in ribosomal RNA (rRNA) maturation (Kiss 2002) and can be found as extracellular RNAs (Kim et al. 2017). piRNAs, initially discovered in germline cells (Klattenhoff and Theurkauf 2008; Girard et al. 2006), are a less studied class of small RNA molecules, however, recent studies have identified them as circulating RNAs (Yuan et al. 2016; Danielson et al. 2017). Besides regulatory sncRNAs, protein-coding mRNAs and tRNAs are also found as circulating RNAs (Yuan et al. 2016) despite their roles in protein synthesis. Furthermore, tRNA-derived small RNAs or tRNA-derived fragments (tRFs) are known to have specific cellular expression patterns (Lee et al. 2009; Zheng et al. 2016) and are associated with some cancer types (Goodarzi et al. 2015). This makes extracellular tRNAs and their fragments potential biomarkers.

Large portions of the human genome are biochemically and transcriptionally active (Pennisi 2012; Djebali et al. 2012; ENCODE Project Consortium 2012). Efforts have been made to deduce the roles of cellular RNAs and their fragments (Palazzo and Lee 2015; Clark et al. 2011; Pauli et al. 2015; Tuck and Tollervey 2011; Scott and Ono 2011; Röther and Meister 2011). The functionality of many extracellular RNA molecules is also an open question (Kim et al. 2017; Yuan et al. 2016), since they can be mere degradation by-products, experimental noise or have alternative roles in circulation.

The aim of this study was to profile RNA molecules in human serum. We analyzed small RNA-seq data from a large (N=477) set of long-term archived serum samples. To assess potential functionality, we analyzed biological variation of sncRNAs and expression/degradation patterns of RNA fragments. To date, this is the most comprehensive analysis of the sncRNA repertoire in human serum.

## RESULTS

### Overall RNA profiles

We analyzed the RNAs in the size range of 17 to 47 nts (Fig 1A). This entails mostly sncRNAs, but it also includes fragments of lncRNAs, mRNAs and other longer transcripts. miRNAs are represented with a peak at 22 nts. The completeness of the profiles relies on sequencing depth, and the saturation analyses showed that canonical miRNAs and tRNAs are approaching plateau with a sequencing depth of about 10-15 Million reads (Fig. 1B). However, the number of piRNAs, isomiRs and tRFs are still increasing at 15 Million reads (Fig. 1B,C).

**Figure 1.**
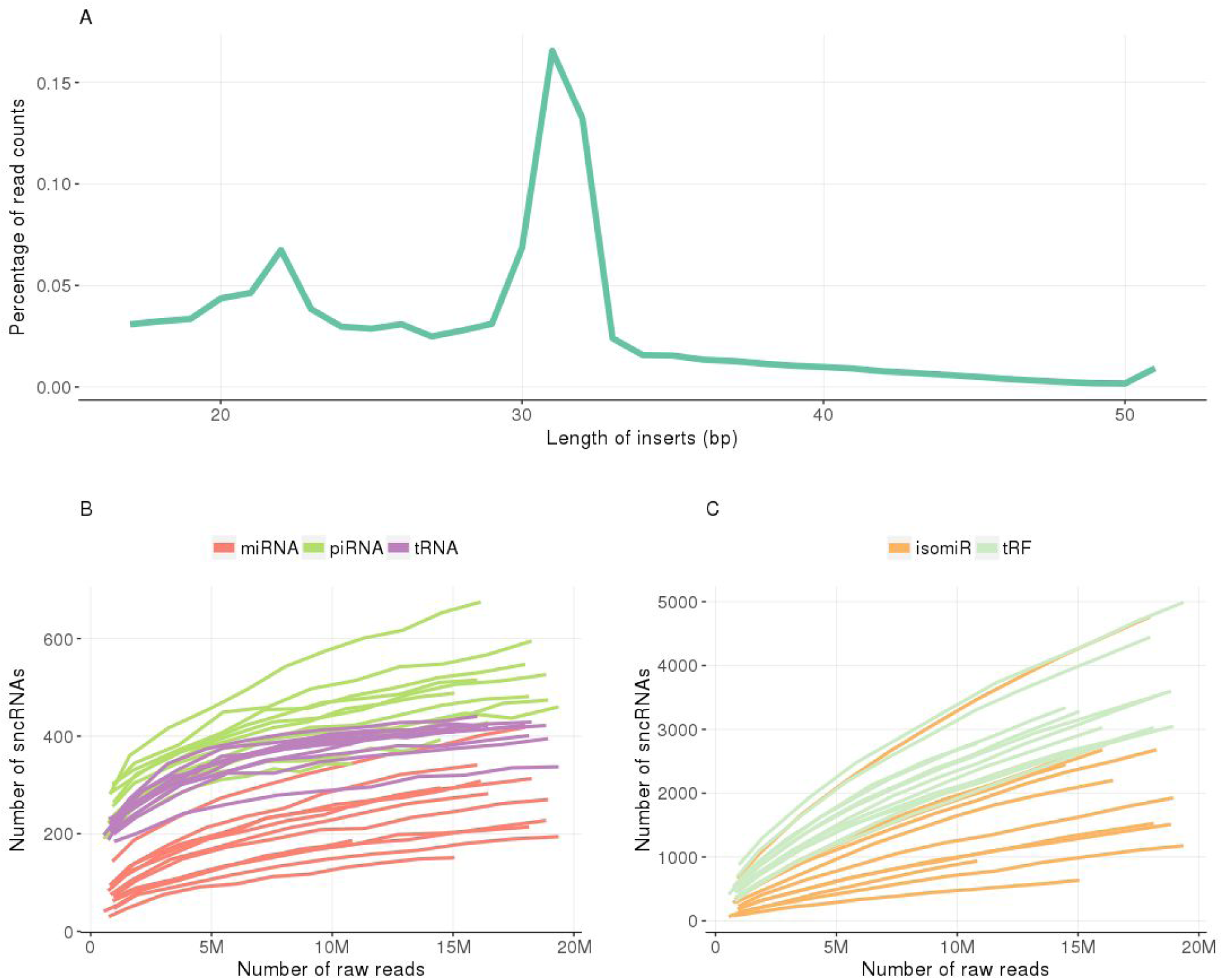
**(A)** The line shows the distribution of trimmed RNA molecule sizes for the serum samples. Our theoretical input library size is between 17 and 47 nts. There are two peaks for the reads at 22 and 31 nts length. This enabled us to detect numerous RNA types including fragments of lncRNAs and mRNAs. **(B)** The saturation lines of canonical genes (i.e. miRNAs, piRNAs, and tRNAs) for a randomly selected subset of serum samples (n=12) are shown. The number of identified genes are still increasing for piRNAs (the dark green lines) but the others are about to reach plateau. **(C)** The non-canonical isoforms (i.e. isomiRs and tRFs) identified are also increasing with the sequencing depth and far from reaching plateau.

We found a total of 258 miRNA, 441 piRNA, 411 tRNA, 24 snoRNA, 125 snRNA and 123 misc-RNA genes that passed the expression threshold that we set (median expression >= 10 reads), representing the core RNA expression profile of serum. In addition, 87 lncRNAs and 1334 mRNAs were detected because of the RNA fragments mapped to these annotations. The transcript origin of RNA reads mapping to multiple genomic locations cannot be determined when mapping qualities are equal for several locations. For comparability to previous studies, we show profiles using both uniquely and multi-mapped reads (Fig. 2A,B). Multi-mapped sequence counts enriches the abundance of high-copy number genes (e.g. piRNA and tRNA). We also used this approach for RNA identification in this study.

**Figure 2.**
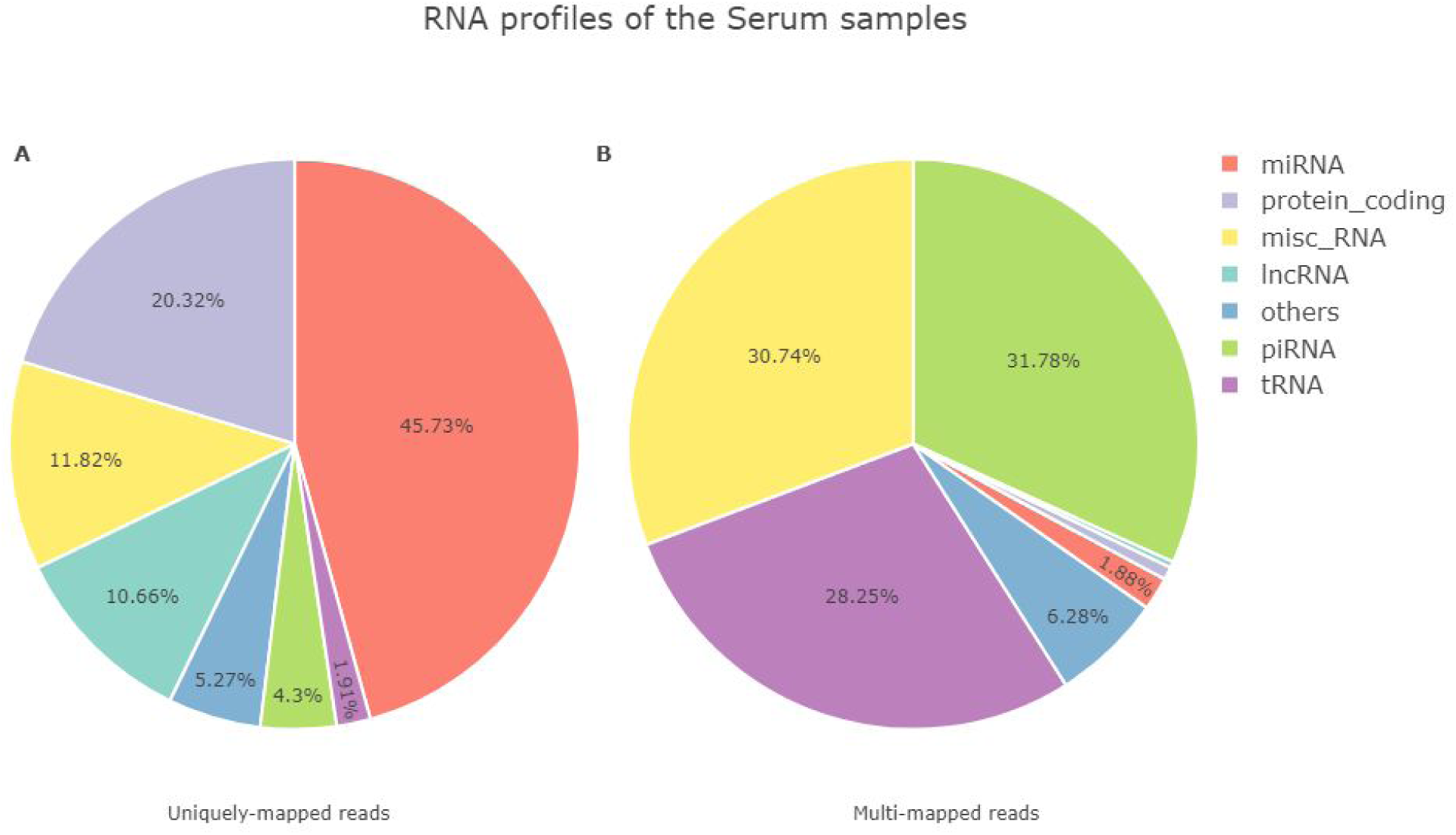
An overall classification of the mapped reads of the serum samples (n=477). **(A)** This pie-chart, generated using uniquely-mapped reads, shows an abundance of miRNA hits followed by protein-coding mRNAs and misc-RNAs. **(B)** Allowing multi-mapped reads is affecting overall RNA profiles. For multi-mapped reads, piRNAs (green) are the most abundant RNA type followed by misc-RNAs (yellow) and tRNAs (purple). The annotations of GENCODE v26 and piRBase were used to create these plots. Similar pie-charts for the technical replicates are at the supplementary (Figure S2).

The overall RNA expression profile shows that some RNA classes are highly expressed compared to others and the top expressed RNAs are listed in Table 1. The misc-RNA class includes Y_RNAs, Signal Recognition Particle (SRP) RNA and Vault RNAs etc. (Table 1). The snoRNAs include U3, U8 and some other related C/D or H/ACA box snoRNAs (Table S4). The snRNAs include U2, U1, U6 and related snRNA genes (Table S5). Complete lists of all identified RNAs are in supplementary tables (Tables S1-S8).

**Table 1.** A summary table of highly expressed RNAs identified in the serum samples.

| Expres<br>sion<br>Rank | miRNA | piRNA | misc-RNA | lncRNA | mRNA |
| --- | --- | --- | --- | --- | --- |
| 1 | hsa-miR-423-5p | piR-hsa-25779 | Y_RNA | RP11-1151B14.3 | NSRP1 |
| 2 | hsa-miR-320a<br>* | piR-hsa-25780 | RNY4** | RP11-20B24.2 | WDR74 |
| 3 | hsa-miR-1246<br>* | piR-hsa-12790 | RNY1** | LINC00910 | VMP1 |
| 4 | hsa-miR-122-5p | piR-hsa-2106 | RN7(x)** | LINC00324 | HOXB4 |
| 5 | hsa-miR-1290<br>* | piR-hsa-25783 | RNY3** | LINC01783 | ATP5G3 |
| 6 | hsa-miR-21-5<br>p | piR-hsa-25782 | SRP | RP11-108M9.3 | MTRNR2L8 |
| 7 | hsa-miR-486-5p | piR-hsa-18709 | VTRNA1(x)** | RP11-473M20.16 | C9orf3 |
| 8 | hsa-miR-148a<br>-3p | piR-hsa-2107 | KCNQ1OT1_5 | CARMN | MTRNR2L12 |
| 9 | hsa-miR-451a | piR-hsa-25781 | 7SK | RNU11 | MTRNR2L1 |
| 10 | hsa-miR-101-3p | piR-hsa-1207 | Vault RNA | RP11-160E2.6 | FAM212A |

Note: \* these miRNAs are challenged, see the Discussion. \*\* similar annotations are collapsed for misc-RNAs. The extended lists are available in Supplementary (Tables S1-S8).

### Isoform profiles of miRNAs and tRNAs

We identified 1642 isomiRs in the serum samples, which passed the detection threshold (i.e. median expression >=10 among samples). The average GC contents of serum isomiRs, canonical forms and miRNA precursors are 0.51, 0.50 and 0.52 respectively. The isomiRs are mostly 3′ isomiRs (78%), followed by 5′ (27%), substitution (22%) and canonical forms (8%). The identified isomiRs are generally an isoform of highly expressed miRNAs (Table 1). For example, hsa-miR-320a, hsa-miR-423-5p, hsa-miR-122-5p and hsa-miR-1246 have 159, 138, 73 and 55 isoforms respectively. A detailed list of the serum isomiRs and their precursors is provided in supplementary (Table S1A).

We identified 1900 tRFs in the serum samples. The average length of these tRNA fragments is ∼29 nts and the average GC content is 0.53. A detailed examination of tRFs showed that they originated from either the 5′ or 3′ end of mature tRNAs (Fig. 3A). This suggests there are no mature tRNAs in serum. The 3′ end of tRNAs was the most abundant region with a uniform distribution throughout a 30 nts region (Fig. 3A).

**Figure 3.**
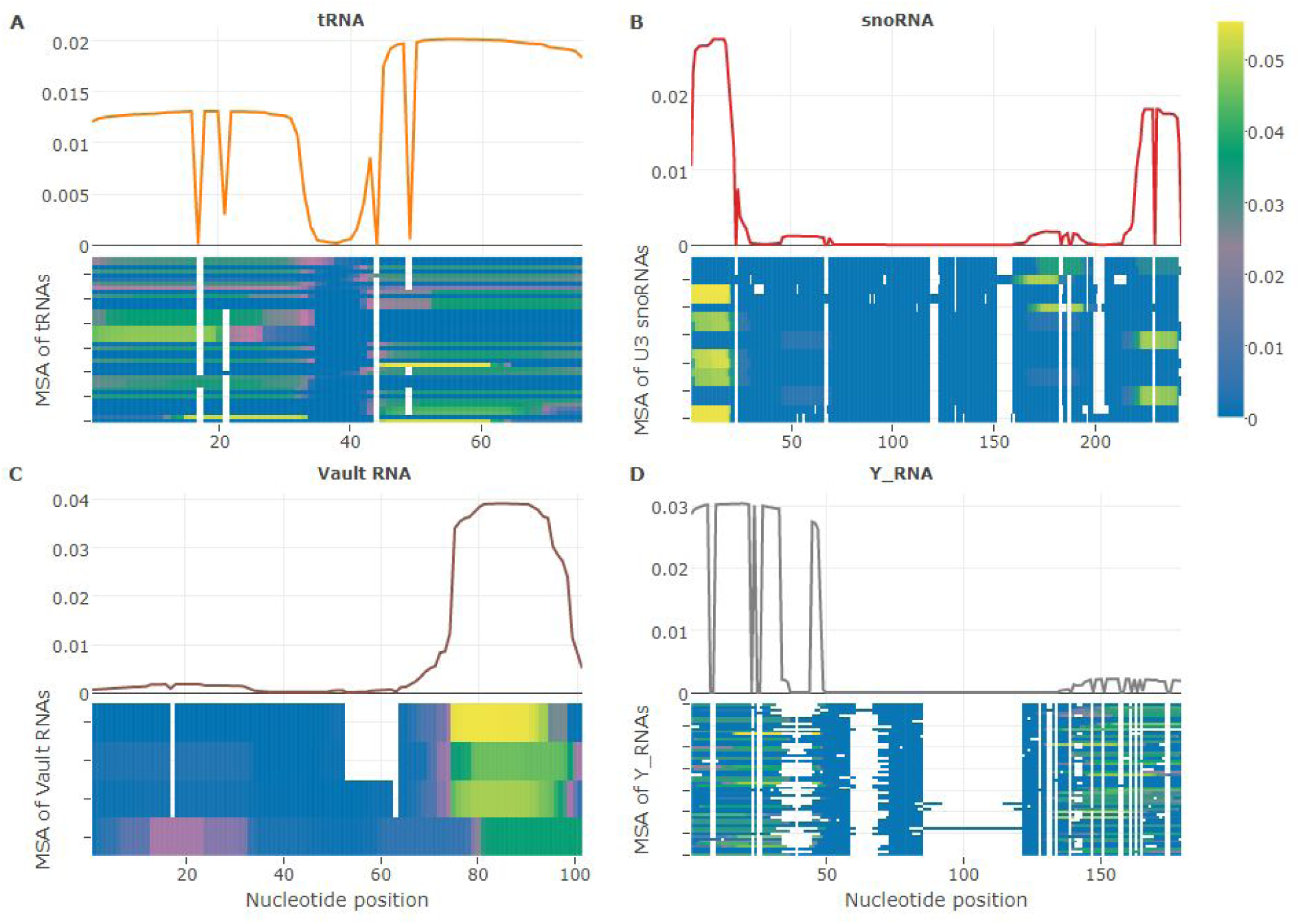
The profiles of mapped reads from highly expressed (A) tRNAs (n=41), (B) U3 snoRNAs (n=18), (C) Vault RNAs (n=4) and (D) Y_RNAs (n=57). Each panel has a multiple sequence alignment (MSA) at the bottom and a corresponding density plot at the top. The x-axes of all plots display a nt position on their MSAs. For example, the MSA of tRNAs is 75 nts long which can be seen at the bottom of the plots. The density plots shows the overall mapping profiles and their x-axes also display nt positions. The heat-maps provide colored representation of the density plot per RNA in the alignment. Yellow and green correspond to the top expressed regions (i.e. high depth), while blue contain almost no mapped reads. White are the gaps in the alignment. **(A)** The reads mapped to mature tRNAs are mostly coming from the 3′ ends (density plot). **(B)** There is a peak at the 5′ end of the snoRNA density plot that corresponds to a 20 nts long region. **(C)** The Vault RNAs identified have a clear signal of expression at their 3′ ends (density plot and yellow bricks at the heatmap). **(D)** The Y_RNA reads are mostly originating from 5′ ends and there is a small peak at the 3′ end (density plot).

### Profiles ofRNA fragments

We also analyzed the profiles of RNA molecules mapped to other annotated regions, including snoRNAs, Vault RNAs, Y_RNAs, mRNAs and lncRNAs. First, U3 snoRNAs are the most abundant within the snoRNA class (Table S4) and the average size of all U3 snoRNA mapped reads is around 29 nts with an average GC content of 0.51. These reads usually come from two regions, the first 20 nts or the last 22 nts region (Fig. 3B), but there are also two smaller peaks between nts 48-74 and 169-195. Second, Vault RNAs have a consistent signal of expression with reads derived from a region covering 75th - 95th nts, while the total size of the Vault MSA is 101 nts (Fig. 3C). These reads also have higher average GC contents, 0.62, than their host Vault RNAs, 0.52. Third, Y_RNAs constitute most of the misc-RNA group’s expression (Table 1). The MSA of Y_RNAs consist of 51 Y_RNAs and 179 nts (Fig. 3D). The expression profiles of Y_RNAs showed that the reads were mapped to the first 1-50 nts region. The average GC content of these reads is 0.51 with an average length of 37 nts. Lastly, as mentioned in the Materials and Methods, we counted the reads only mapped to exonic regions of mRNAs and lncRNAs. The fragments mapped to exonic regions of longer annotations (i.e. mRNA and lncRNA) have average sizes of 29 nts for mRNAs and 30 nts for lncRNAs with GC contents of 0.52 and 0.51 respectively.

### Coefficient of Variation (CV) analyses of sncRNA expression

We analyzed variation in expression of identified sncRNAs to investigate biological signals. In the serum samples, there is a linear relationship between log-normalized mean expression and the standard deviation of identified sncRNAs (Fig. 4A), which shows that the variation is higher for the highly expressed sncRNAs.

**Figure 4.**
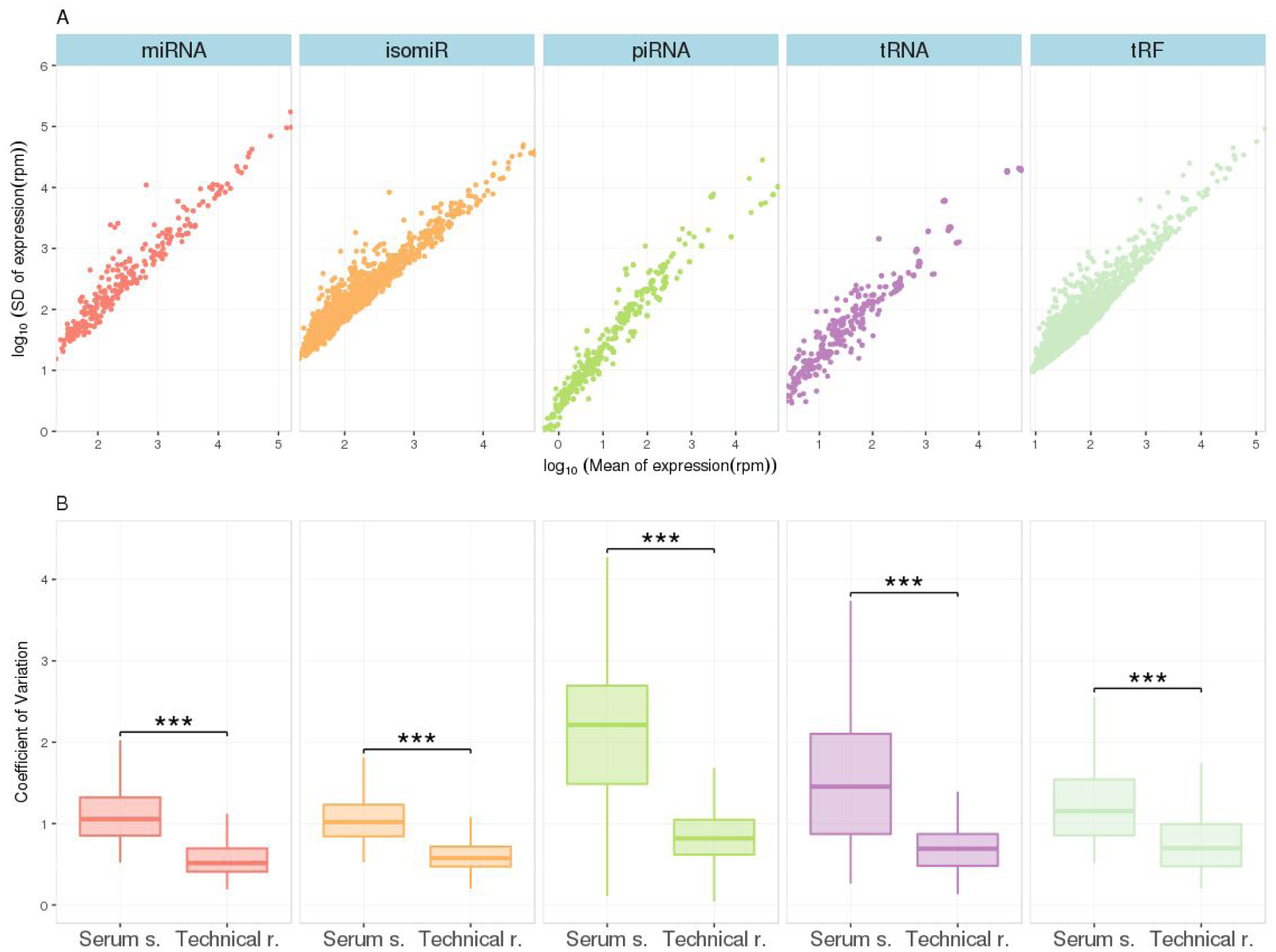
**(A)** The y-axis shows the log_10_ of standard deviations of normalized expression and the x-axis shows the log_10_ mean expression of identified sncRNAs. **(B)** The boxplots show the distribution of CV values in the serum samples and the technical replicates. A pairwise MWU test (*** p ≪ 0.0001) confirmed higher CV values in the serum samples than the technical replicates suggesting higher biological variation for the serum samples than the technical replicates. Randomly generated subsamples of the serum samples (n=17) also produces similar results (Fig. S3) excluding variation due to different samples sizes.

A CV value measures dispersion of a distribution and is a standardised measure of the standard deviation. Distributions of CV values per sncRNA class for both the serum samples and the technical replicates were calculated. We hypothesized that RNA expression in the serum samples will vary more than the technical replicates due to biological variance, because the variation in RNA expression of the serum samples is a combination of technical and biological factors. We tested the null hypothesis: there is no difference in CV values of these two sample sets in three sncRNA types (i.e. miRNA, piRNA and tRNA) and in two different isoforms. We found that the RNA expression varies more in the serum samples than the technical replicates (one sided Mann-Whitney U test (MWU), p ≪ 0.0001 for all) (Fig. 4B). This means that the CV values of RNA expression in the technical replicates are consistently lower than in the serum samples for all sncRNA types, including isoforms (i.e. isomiRs and tRFs).

Low technical variation is preferable for a biomarker (Kahraman et al. 2017), so removing the sncRNAs with high technical variation should create a better set of biomarkers. As an example we tested this with cluster analyses using isomiRs identified both in the serum and technical replicates. The detected isomiRs were divided into four groups based on their CV: all isomiRs (n=1678, identified in both sample groups), low CV (lower than median CV, n=819), very low CV (lower than first quantile, n=40) and high CV isomiRs (higher than median CV, n=859). The four dendrograms created from these groups showed that the low CV and very low CV isomiRs can successfully cluster a set of randomly selected serum samples (n=17) and technical replicates (Fig. S4B,C). However, all isomiRs and the high CV isomiRs cannot successfully cluster these two sample types (Fig. S4A,D). We detected a GC difference between the high CV (0.52) and low CV (0.49) isomiRs (two sided MWU, p=0.003) which may be a reason for the additional technical variation in some isomiRs. Their average internal folding energies, -1.19 kcal/mol for the high and -1.17 kcal/mol for the low CV group, are also slightly different (two sided MWU, p=0.014), which is most likely an effect of the GC difference.

## DISCUSSION

A biomarker is a measurable indicator of a biological state or a phenotype (Lopez et al. 2015; Strimbu and Tavel 2010). There is increasing interest in early-detection of diseases using RNA biomarkers, and numerous studies have investigated circulating miRNAs as candidate non-invasive biomarkers (Rounge et al. 2015; Flatmark et al. 2016; Mitchell et al. 2008; Keller et al. 2011; Maierthaler et al. 2017; Mendell and Olson 2012; Inns and James 2015; Leidinger et al. 2013; Arroyo et al. 2011). We expanded previous research by generating the most comprehensive RNA profile of serum. Our in-depth analyses includes not only miRNAs, but also piRNAs, tRNAs, snoRNAs, snRNAs, misc-RNAs, lncRNAs, mRNAs and RNA fragments such as isomiRs, tRFs, RNA derived particles.

To be able to analyse all the sncRNAs, a size filtering of 15-40 nts is sufficient (Lopez et al. 2015). With our insert size selection (17-47 nts) we were able to do a complete profiling of serum sncRNAs (Fig. 1A). The fragments of lncRNAs, mRNAs and other longer transcripts were also detected in serum. Sequencing depth influences sensitivity of RNA-seq (Fig. 1B) and this is especially notable for isoforms (Fig. 1C). The average sequencing depth is high and selection of a lower threshold (i.e 5) would allow identification of 23% more miRNAs, 10% more piRNAs and 11% more tRNAs, compared to the reported core set (Tables S1-S8). The serum samples in this study can be up to 40 years old, however, the results suggest that many RNA classes are still recoverable with a high expression signal. There is a slight difference between the overall RNA contents of the serum (Fig. 1) and the (fresh) technical replicates (Fig. S2). This difference is mostly likely an artifact of sample pooling rather than of degradation, since the profiles are consistent regardless of sample storage time.

The core set of RNAs were reported by selecting a high expression threshold, which filtered out the RNA products with less stable expression. Our analyses produced comparable results with previous circulating sncRNA profiling attempts. For example, the highly expressed miRNAs in our serum samples, hsa-miR-423-5p, hsa-miR-320a, hsa-miR-122-5p, hsa-miR-486-5p, hsa-miR-486-3p were detected in blood samples (Leidinger et al. 2013; Danielson et al. 2017; Lopez et al. 2015). Hsa-miR-1290 and hsa-miR-1246 were detected in serum and associated with metastasis of lung cancer tumors (Zhang et al. 2016). Some of the highly expressed piRNAs in our serum samples (e.g. piR-hsa-2106 (pir-001311), piR-hsa-27493 (pir-019825), piR-hsa-23209 (pir-020496), piR-hsa-28223 (pir-020388), piR-hsa-28527 (pir-020582), piR-hsa-28374(pir-020485)) are known to exist in plasma and a few of them are also associated with cancer phenotypes (Yuan et al. 2016).

A single miRNA locus can produce various isomiRs with distinct length or sequence (Neilsen et al. 2012) and they have been associated with phenotypes and diseases (Telonis et al. 2017; Morin et al. 2008; Llorens et al. 2013). Both in animal and plants, 3′ isomiRs are the most common ones (Neilsen et al. 2012), consistent with our results. We found that only 8% of the isomiRs are the canonical forms from miRBase, and highly expressed potential isomiRs can be identified in serum. tRFs are other less-known class of sncRNAs which are isoforms of tRNA genes (Lee et al. 2009). They are derived either from mature tRNAs or 3′ of tRNA precursors (Lee et al. 2009; Zheng et al. 2016) and expressed under various stress conditions (Thompson and Parker 2009; Saikia et al. 2014). Many tRFs were associated with different cancer phenotypes (Goodarzi et al. 2015; Zheng et al. 2016) and some were found to be functional like a regulatory miRNA (Maute et al. 2013). Random degradation of tRNAs should give a uniform distribution of tRFs covering the entire tRNA annotation (Zheng et al. 2016). However, we found that tRFs have non-uniform expression patterns (Fig. 3A), suggesting regulated cleavage. This is consistent with known tRF biogenesis (Lee et al. 2009). We also found potentially functional tRNA derived fragments. For example, tRF-5001 was detected in prostate cells in high amounts (Lee et al. 2009). Moreover, 107 tRFs identified were associated with Argonaute family proteins and predicted to have possible mRNA targets (Kumar et al. 2014). One of these 5′ end tRFs have the maximum median expression in our serum samples (Table S3A). It was deposited to MINTbase tRF database (id tRF-30-PNR8YP9LON4V) (Pliatsika et al. 2016) and also found to bind 12 different mRNAs (e.g. EI24, SUGP2 etc.) according to CLASH data (Kumar et al. 2014).

There are RNA fragments originating from well-known annotations, such as snoRNA, misc-RNA, lncRNA and mRNA, that can be functional independent of their host gene (Tuck and Tollervey 2011; Persson et al. 2009; Röther and Meister 2011). In our dataset these RNA fragments are abundant (at least 40% of the all RNA molecules). SnoRNA derived fragments can act like miRNAs to suppress target gene expression (Scott and Ono 2011) and Figure 3B shows that snoRNA in serum also have a non-uniform expression pattern, similar to tRFs. Y_RNAs are short misc-RNAs with functional roles in DNA replication and RNA stability (Kowalski and Krude 2015; Mosig et al. 2007). These fragments, previously found as circulating RNAs in mammals (Nolte-’t Hoen et al. 2012; Kowalski and Krude 2015), have been associated with apoptosis in human cells (Rutjes et al. 1999). Vault RNAs and their fragments were also associated with drug resistance (Persson et al. 2009; Izquierdo et al. 1998). Vault RNAs are a part of ribonucleoprotein complexes (Kedersha et al. 1990; van Zon et al. 2001). They were identified as circulating RNAs in mammals (Nolte-’t Hoen et al. 2012). Both Y_RNAs and Vault RNAs are highly abundant in our serum (Tables 1 and S6) and have a non-uniform expression patterns (Fig. 3C,D). Furthermore, lncRNA and mRNA fragments are known to have different roles such as competing for protein/oligonucleotide binding (Tay et al. 2014; Kulcheski et al. 2016) and target gene regulation (Pircher et al. 2014; Rogler et al. 2014). The RNA fragments mapped to them have similar size and GC distribution with other sncRNA fragments in our dataset. The expression is often high and stable for these fragments and they cover only small fractions of their host gene (i.e. non-uniformity).

An important question is whether the discovered sncRNAs and their fragments are genuine functional products. The above mentioned high expression pattern and regulated cleavage suggest function. Random degradation and experimental noise from RNA-seq studies (McIntyre et al. 2011; Backes et al. 2016; Tarazona et al. 2011; Marioni et al. 2008) might introduce false positive prediction of biological function and associations due to lack of RNA-seq sensitivity (Todd et al. 2016; McIntyre et al. 2011). We proposed that CV analysis (Fig. 3 and Fig. S4) is suited for suggesting biological variation, because in an ideal setting, technical replicates should contain no biological variation, only technical variation. However, variation in serum samples is a sum of both biological and technical variability. We identified a statistically significant difference in average CV between technical and serum samples for all sncRNA classes (including isoforms) that shows higher variation for serum samples. This supports a biological signal in serum RNA expression and suggests potential function for circulating RNA molecules.

Technical variation in RNA-seq may vary depending on RNA molecule characteristics such as expression level, size, sequence and secondary structure. We measured a range of CV values in our technical replicates even though we expected them to be closer to zero (Fig. 4B). High technical variation can decrease biomarker value by influencing reproducibility. This can be observed in our cluster analysis: the low CV and very low CV isomiRs best discriminate the serum and technical replicate group. We detected a statistically significant difference between the GC contents of high and low CV isomiRs which may partly explain technical variation. Some of those highly discriminatory isomiRs (e.g. isomiRs of hsa-miR-192-5p, hsa-miR-375 etc.) were successfully clustering various cancer tissues in a binary classification approach (Telonis et al. 2017). Another 5′ end isoform of hsa-miR-101-3p, with a low technical variation in our study, was also found to have a role in gene silencing in brain tissues (Llorens et al. 2013). In short, this analysis showed that a set of isomiRs with low CV is less prone to technical variation and they successfully cluster the two groups.

The large sample size, high coverage and the diversity of RNA products analyzed are the strengths of our study. We extensively profiled abundant RNA fragments in serum, and showed specific cleavage patterns of some RNA fragments for the first time. We also utilized a set of technical replicates to measure biological signal of serum RNA expression. This analysis suggested functionality for RNA fragments. However, there are potential limitations that we should address.

First, long-term storage may degrade some unstable RNAs, though our results suggested that the degradation is not substantial for sncRNAs. It has been proven for miRNAs that they remain stable in severe conditions (Chen et al. 2008) and in circulation (Arroyo et al. 2011). They can be extracted from long-term serum (Zhu et al. 2009; Rounge et al. 2015). Moreover, any RNA found in serum stored up to 40 years is evidently quite stable, which is one of the critical criteria for good biomarkers. Second, although all samples are processed in the same way, slight differences in laboratory processing may still introduce some technical variance into expression which cannot be removed totally. We addressed this variation (Fig. 4B) using the technical replicate samples and CV values, which showed higher technical variation was introduced into some sncRNAs than the others. Third, the lab and bioinformatic analysis methods chosen may compromise generalizability of results. For example, differences in gel cut size will change proportions of sncRNAs and narrower cut will limit detection of certain sncRNA classes. Detection threshold and allowing multi-mapped reads will also change the overall RNA profiles substantially (Fig. 2). Selection of read mapper and algorithm parameters are other bioinformatics related factors that can influence overall results (Ziemann et al. 2016). Furthermore, high quality annotations are also essential to correctly identify transcripts (Harrow et al. 2012), which is still a major barrier even for well-studied human miRNAs (Fromm et al. 2015). For example, highly expressed miRNAs, hsa-miR-1246 and hsa-miR-320a, are questioned for not being a miRNA gene (Fromm et al. 2015). Since they are part of miRBase, we reported them (and their isoforms) as miRNAs to be consistent with the literature. However, improving annotation quality is an on-going process and still far from perfect. It is also reasonable to consider possible alternative functions of the RNA fragments derived from longer host genes rather than counting them as a single piece of a large annotation. For instance, counting tRFs or misc-RNA derived fragments as their host genes would have overshadowed the specific expression patterns that we reported in Figure 3.

## CONCLUSION

Here we present a comprehensive characterization of human serum sncRNA content. Our results unveiled that most of the RNAs identified in serum are not random by-products but most likely have roles as circulating RNAs. This conclusion is supported by (1) stable high expression, (2) biological signal and (3) distinct expression patterns of many identified RNA molecules. Our results suggest new opportunities for novel biomarker discovery in serum, but they are also transferable to other body fluids and tissues.

## MATERIALS AND METHODS

### Study design

The JSB cohort is a population-based cancer research biobank containing pre-diagnostic serum samples from 318 628 Norwegians (Langseth et al. 2016). By linking data from the Cancer Registry of Norway (Larsen et al. 2009) with the JSB cohort, we identified serum donors (n= 477) that were cancer-free at least 10 years after sample collection. We do not have any information about non-malignant conditions. A previous study showed that miRNA (and other sncRNA) discovery is possible in long-term archived serum samples (Rounge et al. 2015). In addition to investigate technical variation, fresh serum from 6 individuals were pooled into one sample and divided into 17 aliquots. They were analysed as technical replicate samples. The downstream analyses were identical for all samples (Fig. S1). The donors have given broad consent for the use of the samples in cancer research. The study was approved by the Norwegian regional committee for medical and health research ethics (REC no: 2016/1290).

### Laboratory processing

RNA was extracted from 2 x 200 μl serum using phenol-chloroform phase separation and the miRNeasy Serum/Plasma kit (Cat. no 1071073, Qiagen) on a QIAcube (Qiagen). Glycogen (Cat. no AM9510, Invitrogen) was used as carrier during the RNA extraction step. Small RNA-seq was performed using NEBNext^®^ Small RNA Library Prep Set for Illumina (Cat. No E7300, New England Biolabs Inc.). Size selection was performed using a 3% Agarose Gel Cassette (Cat. No CSD3010) on a Pippin Prep (Sage Science) with a cut size optimized to cover RNA molecules from 17 to 47 nt in length. Sequencing libraries were indexed and 12 samples were sequenced per lane of a HiSeq 2500 (Illumina).

### Bioinformatics analyses

The total number of reads generated was approximately 10 billion. The average sampling depths of the serum and technical replicate samples were 17.9 and 19.5 million raw reads, respectively. The reads were initially trimmed for adapters using AdapterRemoval v2.1.7 (Schubert et al. 2016). We then mapped the collapsed reads (generated by FASTX v0.14) to the human genome (hg38) using Bowtie2 v2.2.9 (10 alignments per read were allowed). We compiled a comprehensive annotation set from miRBase/MirGeneDB (Kozomara and Griffiths-Jones 2014; Fromm et al. 2015) for miRNAs, pirBAse/pirnabank for piRNAs (Zhang et al. 2014; Sai Lakshmi and Agrawal 2008), GENCODE (Harrow et al. 2012) for other RNAs and tRNAs. We used SeqBuster (Pantano et al. 2010) to get isomiR and miRNA profiles of our samples. To count the reads mapped on other RNAs, HTSeq (Anders et al. 2014) was utilized in a Python script. We used a threshold of 10 median read count per sncRNA to get a robust signal of expression. For longer transcripts (e.g. messenger RNA (mRNA) or long non-coding RNA (lncRNA)), we counted reads only mapped to exonic regions. However, this does not mean that the non-exonic mapped reads are not important. We are interested in bona fide fragments of longer genes but many non-exonic reads usually overlap with other short annotations, so it can be hard to determine their correct origin. Read counts were normalized to get reads per million (RPM) values. The coefficient of variation (CV) was calculated based on RPM values for the genes identified both in the serum and technical replicates in order to test biological and technical variation.

In order to get isoform and coverage profiles of tRNAs, we counted the reads mapped to tRNAs. There are 649 mature tRNA annotations available in GENCODE. We selected 41 tRNAs accounting for 99% of all reads mapped to tRNA annotations. The tRNAs were aligned to Rfam model (RF00005) using the *cmalign* tool (Nawrocki and Eddy 2013) to get a multiple sequence alignment (MSA) of expressed tRNAs. Similar analyses were conducted for U3 snoRNAs and other miscellaneous RNAs (misc-RNA) (the models are RF00012, RF00006 and RF00019). Misc-RNAs denote RNA transcripts that are not classified into any other groups (Harrow et al. 2012), which were taken from Rfam (Nawrocki et al. 2015).

## Acknowledgements

We would like to acknowledge Tove Slyngstad and Kristina Kymre for performing lab and coordination tasks. The sequencing service was provided by the Norwegian Sequencing Centre (www.sequencing.uio.no), a national technology platform hosted by Oslo University Hospital and the University of Oslo supported by the “Functional Genomics” and “Infrastructure” programs of the Research Council of Norway and the Southeastern Regional Health Authorities.

## DATA AVAILABILITY

Data is available with restricted access at European Genome-phenome Archive (EGA) accession number xx-xx.

